# Selecting sites for strategic surveillance of zoonotic pathogens: a case study in Panamá

**DOI:** 10.1101/2024.08.12.607103

**Authors:** Marlon E. Cobos, Jonathan L. Dunnum, Blas Armién, Publio González, Enós Juárez, Jacqueline R. Salazar, Joseph A. Cook, Jocelyn P. Colella

## Abstract

Surveillance and monitoring of zoonotic pathogens is key to identifying and mitigating emerging public health threats. Surveillance is often designed to be taxonomically targeted or systematically dispersed across geography, however, those approaches may not represent the breadth of environments inhabited by a host, vector, or pathogen, leaving significant gaps in our understanding of pathogen dynamics in their natural reservoirs and environments. As a case study on the design of pathogen surveillance programs, we assess how well 20 years of small mammal surveys in Panamá have sampled available environments and propose a multistep approach to selecting survey localities in the future. We use >8,000 georeferenced mammal specimen records, collected as part of a long-term hantavirus surveillance program, to test the completeness of country-wide environmental sampling. Despite 20 years of surveillance, our analyses identified a few key environmental sampling gaps. To refine surveillance strategies, we selected a series of core historically sampled localities, supplemented with additional environmentally distinct sites to more completely represent Panama’s environments. Based on lessons learned through decades of surveillance, we propose a series of recommendations to improve strategic sampling for zoonotic pathogen surveillance.

## Introduction

With the majority of emerging infectious diseases jumping to humans and domestic livestock from wildlife, scientists and society are grappling with how to effectively monitor zoonotic pathogens and their wild hosts. Traditional surveillance efforts often target specific taxa—for example, a few target species of bats or rats—or localities of immediate interest, often near outbreak sites. Broader attempts at wildlife surveillance have prioritized geographically dispersed sampling, however, those strategies often do not fully characterize the suite of available environments in a region, resulting in an incomplete representation of existing diversity (Hortal and Lobo 2005; Hortal et al. 2015), that includes reservoir host, vector, and pathogen species. Common environments are likely to be overrepresented by such a geographically dispersed sampling approach, whereas rare environments or community assemblages may be missed altogether (Nuñez-Penichet et al. 2022). Explicit consideration of environmental conditions, in addition to geography, may better guide efficient biodiversity inventories and monitoring (Velásquez-Tibatá 2019; D’Antraccoli et al. 2020). For zoonotic pathogens, and particularly those strongly influenced by environmental conditions (e.g., hantaviruses and Rift Valley Fever; Yates et al. 2002; Nanyingi et al. 2015), sampling host diversity across the breadth of available environments is important for understanding enzootic transmission cycles, identifying conditions associated with spillover, and ultimately, to proactively prevent loss of human life and damage to local economies and livelihoods.

Zoonotic pathogen surveillance has historically been reactive. That is, surveys are planned and executed after a public health threat has emerged (Vrbova et al. 2010). An example of such surveys followed the 1993 outbreak of *Sin Nombre* hantavirus (SNV) in the American Southwest. Hantaviruses (Family: Orthohantaviridae) typically can cause mild to severe hantavirus pulmonary syndrome (HPS) in the Americas and hemorrhagic fever with renal syndrome (HFRS) in Eurasia (McCaughey and Hart 2000). To identify the causal agent for a series of mysterious deaths in the four-corners region of the United States, the Museum of Southwestern Biology (in collaboration with the Centers for Disease Control and Prevention, New Mexico Department of Health, and the Indian Health Service) conducted holistic small mammal surveillance resulting in thousands of physical specimens and associated frozen tissues. That outbreak marked the beginning of integrated collaborations between public health agencies, virologists, ecologists, and museum scientists that completely reshaped our understanding of hantavirus systematics, evolution, and ecology (Mills et al. 1999; Dunnum et al. 2017).

Those resources were used to identify *Peromyscus* rodents (deer mice) as the primary reservoir, of *Sin Nombre* hantavirus, with emergence shown to be closely linked to El Nino cycles and related environmental variation (Stenseth et al. 2002; Yates et al. 2002; Guterres and Sampaio de Lemos 2018). Because most zoonotic pathogens exhibit density-dependent transmission, infection risk for rodents increases at higher population densities (Anderson and May 1979; Wilson et al. 2002). Therefore, understanding the connection between pathogen, host, and environmental variables provided critical public health guidance, both proactively (e.g., prediction of deer mouse population explosion; Glass et al. 2006); and retroactively (e.g., risk reduction through proper cleaning of rodent excrement; Escobar and Morand 2021), further reinforcing the importance of voucher-backed wildlife surveillance.

Proactive zoonotic pathogen surveillance and discovery have gained more interest after the global COVID-19 pandemic (Leifels et al. 2022), also caused by a zoonotic pathogen originally circulating in Southeast Asian mammals (Perveen et al. 2021; Fischhoff et al. 2021). Such surveillance initiatives aim to characterize baseline pathogen prevalence and distribution in native, wild reservoirs prior to disease emergence in humans (Daszak et al. 2020; Colella et al. 2021). To completely characterize zoonotic pathogen diversity in an area (e.g., region, state, country), we must consider both what is known and what has not yet been explored (see similar ideas for biodiversity inventory in Medina et al. 2013; Nuñez-Penichet et al. 2022). For example, historical sampling, distribution of novel environments, proximity to outbreak sites and human habitation, among other relevant details provide key *a priori* information that can guide site selection. Longitudinal information (e.g., multi-decadal sampling) is further critical to understanding the natural enzootic dynamics, probability of pathogen transfer to humans or livestock, and complex mechanisms of disease outbreak and transmission (Brouwer et al. 2022). Beyond known information, however, many geographic locations and distinct environments remain unsampled. Among macrofauna, the strong relationship between species richness and environment theoretically also holds true for microfauna, like parasites and pathogens. Therefore, sampling a breadth of distinct environments is expected to saturate pathogen species accumulation curves more rapidly than simply geographically dispersed sampling.

Repeated spillover of hantaviruses from wildlife into humans in the last few decades, combined with associated high mortality rates for some of these viruses (Dearing and Dizney 2010), has increased public concern and awareness of the disease (Grange et al. 2021). For example, in the early 2000s, spillover of *Orthohantavirus chocloense* (CHOV) into humans occurred on the Azuero Peninsula of Panamá (Bayard et al. 2004). Serological surveys of household members and neighbors of those infected found antibody prevalences around 13% (Bayard et al. 2004; Kuhn et al. 2023). Serological tests of wild mammals identified *Oligoryzomys costaricensis,* the Costa Rican pygmy rice rat, and *Zygodontomys brevicauda*, the short-tailed cane mouse, as the primary reservoir hosts of Choclo and Calabazo hantavirus strains, respectively (Vincent et al. 2000; Bayard et al. 2004). The outbreak notably resulted in the cancellation of Carnival, a major Panamanian festival, which had cascading negative effects on local economies. In response, the Gorgas Memorial Institute for Health Studies partnered with the Museum of Southwestern Biology at the University of New Mexico, the same biorepository involved in 1993 hantavirus surveillance in the U.S., to implement a country-wide small mammal monitoring program (Gonzalez et al. 2023). The surveillance program was designed to be geographically dispersed, with sites in 12 of 13 Panamanian Provinces (i.e., Provinces and Comarcas), and environmentally broad, representing seven of nine ecoregions in the country. There are now, however, higher resolution tools for quantitatively evaluating environmental variation that can be used to refine current surveillance strategies. Sampling in Panamá occurred throughout the year (i.e., rainy and dry seasons) to capture seasonal and temporal variation, which are known to affect both the host and pathogen (Mills and Childs 1998; Armién et al. 2016). Those surveys spanned 2000–present and resulted in the permanent archival of >8,000 holistically sampled mammal specimens. Collected specimens were screened for the presence of hantavirus antibodies, and preliminary explorations of those data suggest significant relationships between aspects of seasonality and hantavirus infection dynamics in wild rodents (Gonzalez et al. 2023).

Physical specimens yielded from hantavirus surveillance in Panamá have helped identify a tentative link between hantavirus emergence in humans and environmental variation, but this multidecadal effort has also proven economically and logistically challenging. Lessons learned during this long term surveillance program now provide an opportunity for review with the aim of adaptively improving sampling strategies. Our main goals with this study are to assess the completeness of environmental sampling in Panamá following 20 years of hantavirus surveillance and to select optimal sites for zoonotic pathogen surveillance moving forward, considering historical data and environmental and geographic factors in Panamá. We identified key sites for continued surveillance and potential future expansion. We used hantaviruses in Panamá as a case study due to the depth and breadth of publicly available, standardized, machine-readable, and voucher-backed surveillance data; however, the surveillance strategies outlined here are relevant to diverse zoonotic pathogens, hosts, and geographies.

## Methods

### Summary

We outline a multistep approach to select strategic survey sites based on ∼20 years of small mammal surveillance in Panamá led by the Gorgas Memorial Institute for Health Studies (Panamá) in collaboration with the University of New Mexico’s Museum of Southwestern Biology. Historical specimen collection localities were combined with biogeographic, environmental, and socio-economic factors for site selection (Fig. 1). We assessed the completeness of historical sampling of available environmental and geographic conditions in the country. We then identified a reduced set of historical sampling locations that allowed us to cover strategic geographic areas in the country and, perhaps more importantly, most environments. After that, we identified unsampled environmental conditions in accessible areas in the country and selected additional, accessible sites for sampling to complete our previous selection. Selected sites were vetted by in-country stakeholders engaged in the original surveys, regarding accessibility, before considering them as part of an updated survey strategy. All analyses performed herein can be reproduced following the methods and using R code provided in a GitHub repository at https://github.com/marlonecobos/Hanta_PAN_suitability.

**Figure 1.**
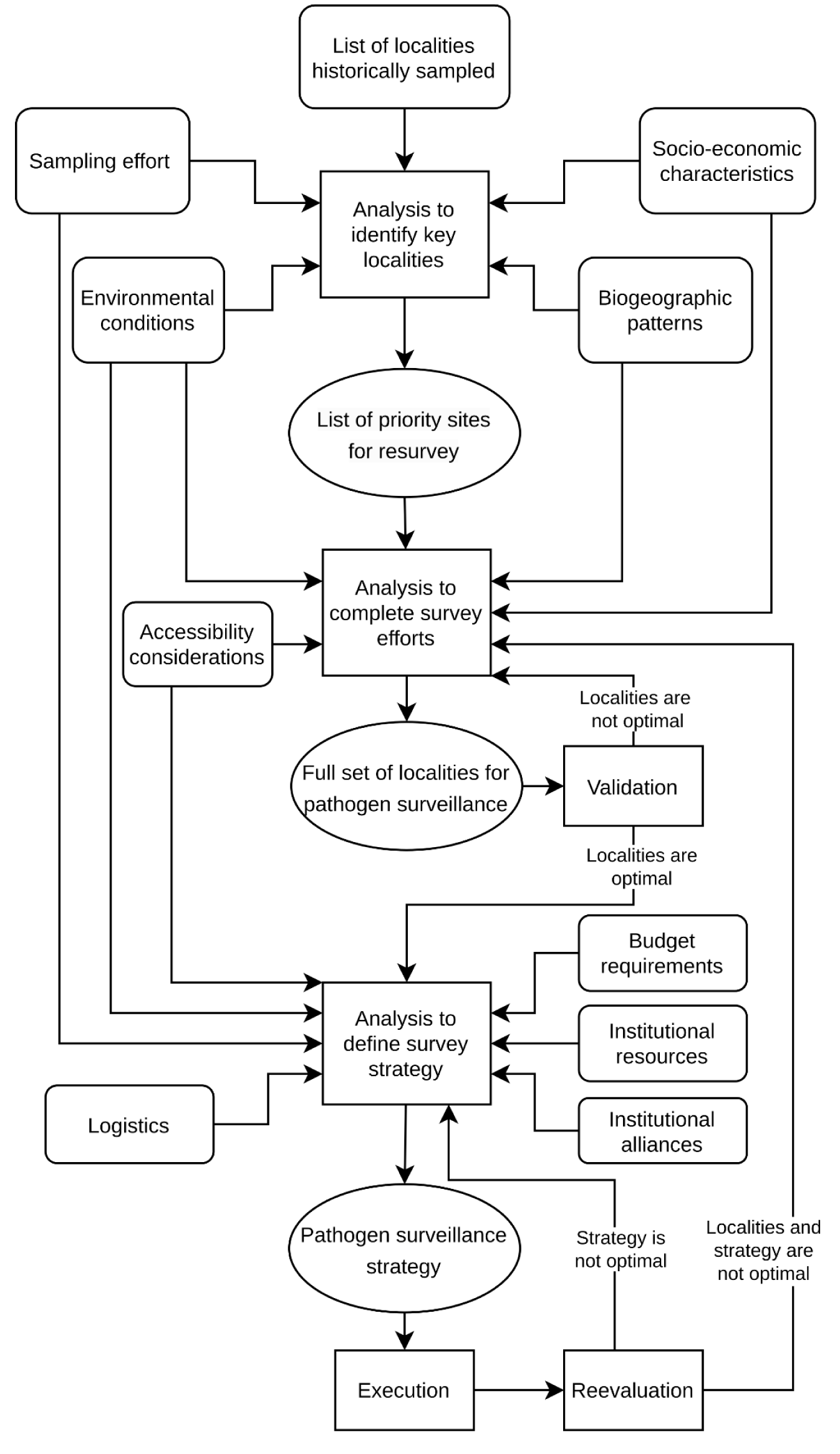
Schematic workflow of the process used to identify sites for surveillance of zoonotic pathogens.

### Data

We downloaded all records of mammals from Panamá that have been serologically screened for Hantavirus (Gonzalez et al. 2023) from the Arctos museum database (https://arctos.database.museum; Cicero et al. 2024). Although our goal is to refine sites for general zoonotic pathogen surveillance, these data represent the most comprehensive pathogen survey effort in the country and are unprecedented for both species diversity and temporal scale. For this reason, our analysis used these specimen records as a starting point to understand how well historical surveys captured environmental variation in Panamá. The search criteria were as follows: Collection = “MSB:Mamm”, Any Geographic Element = “Panama”, date range = “2000–2020”, Class = “Mammalia”, Has Tissue = “yes”. The data export contained the specific collection locality (geographic coordinates), host species identity, hantavirus serological test-result (positive/negative), and the date of collection, among other variables. Specimen records without geographic coordinates or serological results were not considered.

We obtained spatial polygons of the administrative borders of Panamá and its provinces from the GADM database (www.gadm.org). We downloaded spatial layers of roads in the country from the STRI GIS Data Portal (https://stridata-si.opendata.arcgis.com), and created a 2.5 km buffer around all roads to represent areas accessible via ground transportation. Biogeographic aspects of the country were considered in our explorations using the land use cover layer for Panama (Ministerio de Ambiente 2017), and the Köppen-Geiger climate classification maps (Beck et al. 2018). To consider the hantavirus cases in humans, we used the information presented in Armién et al. (2023).

To represent environmental conditions in Panamá, we used variables from the Chelsa-climate database at a spatial resolution of 30 arc-seconds (∼1 km; available at https://chelsa-climate.org; Karger et al. 2017). We used the variables annual mean temperature (bio1), maximum temperature of warmest month (bio5), minimum temperature of colder month (bio6), temperature annual range (bio7), annual precipitation (bio12), precipitation seasonality (bio15), precipitation of wettest quarter (bio16), precipitation of driest quarter (bio17), maximum potential evapotranspiration, mean potential evapotranspiration, minimum potential evapotranspiration, maximum vapor pressure deficit, mean vapor pressure deficit, and minimum vapor pressure deficit. We masked these layers to the administrative area of Panamá. To summarize environmental dimensions into fewer variables, we performed a principal component analysis (PCA) with the layers. We used the first two principal components (PCs) in further analyses, as they summarized most of the variance from all variables. We then masked the PCs to the areas accessible in the country (within the 2.5 km road buffer) to characterize the suite of environmental conditions present in areas that could be realistically sampled. We further extracted environmental information for the collection localities and dates associated with each specimen record to characterize the subset of sampled environmental conditions.

We downloaded data from the GADM database using the package *geodata* (Hijmans et al. 2023) in R v. 4.2.2 (R Core Team 2023). Spatial vector and raster processing was done using the R package *terra* (Hijmans 2023) and QGIS v3.22 (QGIS Development Team 2019).

### Historical sampling and accessibility

To understand how historical hantavirus surveillance was done within the geographic limits of Panamá, we used raster cells at ∼1 km resolution. These cells hereafter represent candidate localities for resurvey and analysis. We quantified the number of sampling days, number of species, number of specimens collected, and hantavirus seroprevalence per each 1 km cell (i.e., locality) across the country. Localities with higher values for all of these quantifications were highlighted for further analysis, as they can be important for future longitudinal characterizations of zoonotic pathogen communities.

We explored historical sampling in terms of sampled environmental conditions by visually comparing the cloud of environments present in Panamá against the values of sampled environments. We calculated a kernel-density for environmental conditions in Panamá and in accessible areas. We also identified unsampled environmental conditions regardless of accessibility. To quantify differences between sampled environments and the entire set of conditions in Panamá, we used the mobility oriented-parity metric (MOP; Owens et al. 2013). This analysis maps levels of environmental dissimilarity (distance) between a set of reference points (sampled localities) and the area of interest (Panamá). We also used the MOP analysis to compare environments in accessible areas against those in the whole country to identify how different environments are in the country compared to those in accessible areas. Kernel density calculations were performed using the R package *ks* (Duong 2022). MOP analyses were performed using the *mop* package (Cobos et al. 2024). All other analyses were performed with base R functions.

### Selection of core historical sites for continued surveillance

From among historically sampled localities, we selected a set of “core” sites for zoonotic pathogen surveillance in Panamá. To make that selection, we considered the following criteria in concert: (1) sites with distinct environmental conditions, to “evenly” cover the breadth of available and sampled environments in Panamá (our main criterion); (2) sites with high sampling effort; (3) sites of relevance for hantavirus monitoring based on the number of specimens collected and hantavirus seroprevalence; (4) sites that represent the major biogeographic regions in Panamá; and (5) sites that are close to human settlements or related to human activities that can generate social conflicts (e.g., migratory movements in the Darién region). All core sites are accessible, as they are localities that have been historically sampled.

To explore interactively the relationship of candidate cell localities with other geographic features (roads, human settlements, land use, etc.), we used Google Earth Pro (Google 2024; https://www.google.com/earth/). This interactive exploration allowed us to pick cell localities that are accessible but also meet most of the criteria described above.

### Additional strategic sites for surveillance

To supplement the core set of historically surveyed sites, we identified additional, unsampled localities that could improve the coverage and representation of novel environmental conditions in accessible areas. To explore environmental conditions in a regionalized manner, as opposed to point-by-point, we divided the environmental space in Panamá (Fig. S1) into a grid of 20 rows and 20 columns, resulting in ∼400 environmental blocks, each containing a collection of points that are environmentally similar (Nuñez-Penichet et al. 2022). Then, we selected blocks from within clusters of unsampled environmental cells that were maximally distanced from core sites to sampled environments more uniformly. If historically sampled localities were available in the selected blocks, we picked one locality per block, considering the remaining criteria for site selection. If no historical localities existed within the selected block, we chose a new locality trying to meet most of the aforementioned criteria. We performed this environmental data processing and exploration step using the R package *biosurvey* (Nuñez-Penichet et al. 2022). We used the package *terra,* base R functions, and interactive explorations in Google Earth Pro to complete site selection (Fig.S1).

## Results

### Data processing results

Our downloaded dataset included 8,090 georeferenced and serologically screened specimen records (dated 2000-2018) from 36 identified mammal species, including three non-natives: *Mus musculus, Rattus norvegicus*, and *Rattus rattus* (Table S1). Sixteen specimens were not identified to species level. The species with the most specimen records were, *Zygodontomys brevicauda* (*n* = 3,513), *Sigmodon hirsutus* (1,186), and *Liomys adspersus* (886). *Oligoryzomys costaricensis*, the known host of *Orthohantavirus chocloense*, had a total of 822 records. The species with the highest hantavirus serological prevalence were *Reithrodontomys sumichrasti* (0.44), *Reithrodontomys mexicanus* (0.29), and *Oligoryzomys costaricensis* (0.17), but antibodies were detected in 10 species.

The first two PCs derived from environmental variables summarized 75% of the variance (Table S2). The first PC was highly loaded by mean and maximum vapor pressure deficit, as well as maximum temperature of the warmest month (Table S3). The second PC was loaded mainly by temperature annual range and precipitation seasonality.

### Historical sampling in Panamá

Localities sampled are strongly associated with areas that are accessible by roads (Fig. 2). Although the breadth of environments in Panamá are fairly well represented within 2.5 km of a road, not all of those environments are yet represented by historical sampling. Sampling effort was highest in the provinces of Chiriquí and Los Santos, with species diversity highest in Chiriquí, Los Santos, Darién, and Coclé (Fig. 2). Most seropositive rodents were detected in the Los Santos province.

**Figure 2.**
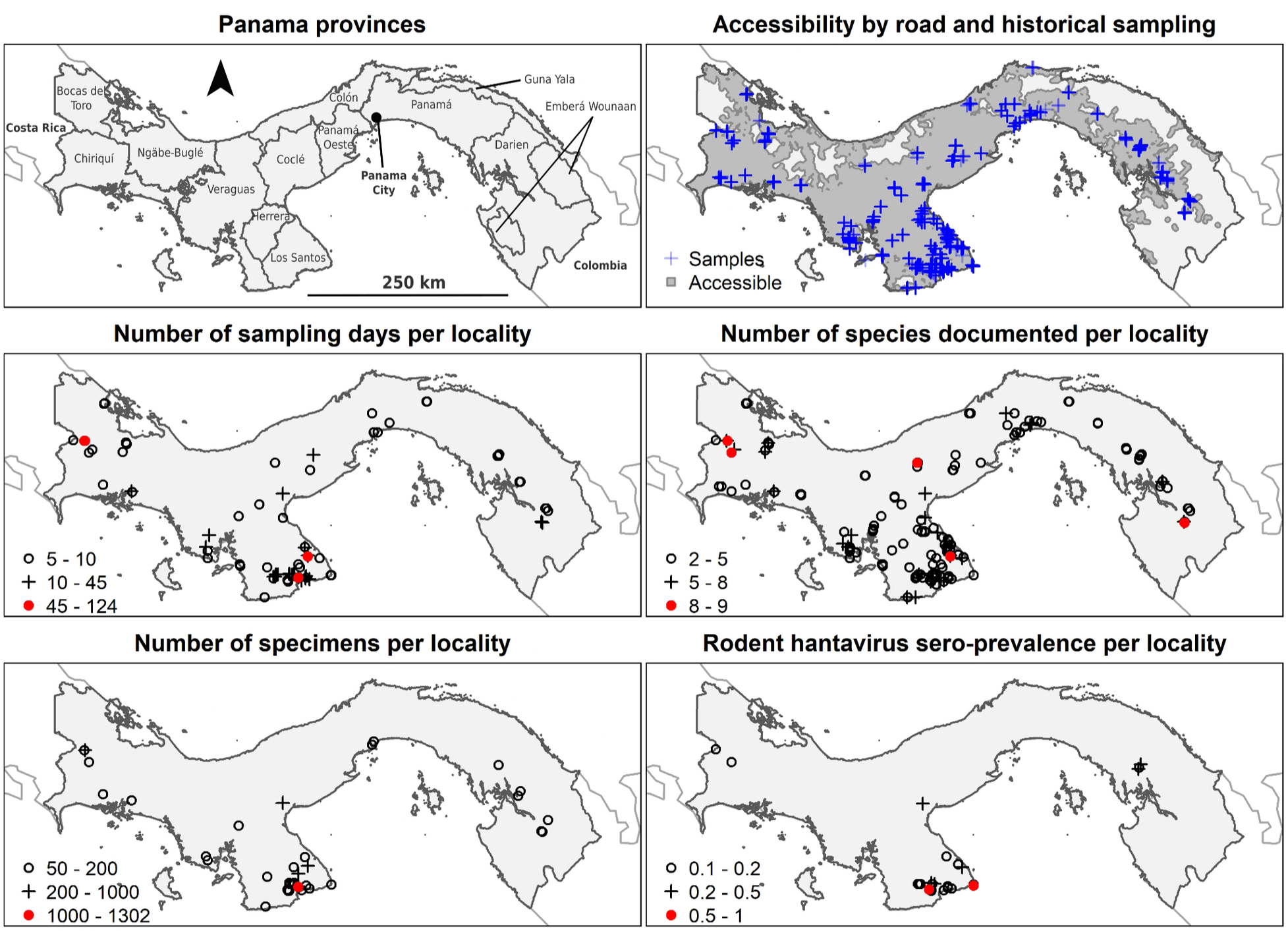
Summary of historical sampling and road accessibility in Panamá. Samples refer to records of collected mammal specimens that have been serologically screened for hantavirus. Localities with lower values in terms of sampling days, species, specimens, and seroprevalence are not shown to help visualize higher priority localities.

Geographic sampling gaps are clustered in areas that are inaccessible or have limited accessibility (Fig. 3). MOP analyses showed two main patterns: (1) the environments represented in historical sampling are comparable with conditions in most parts of the country, with the exception of areas in the Northwest; and (2) most of the environmental conditions in the country are represented in areas that are accessible by road (Fig. 3).

**Figure 3.**
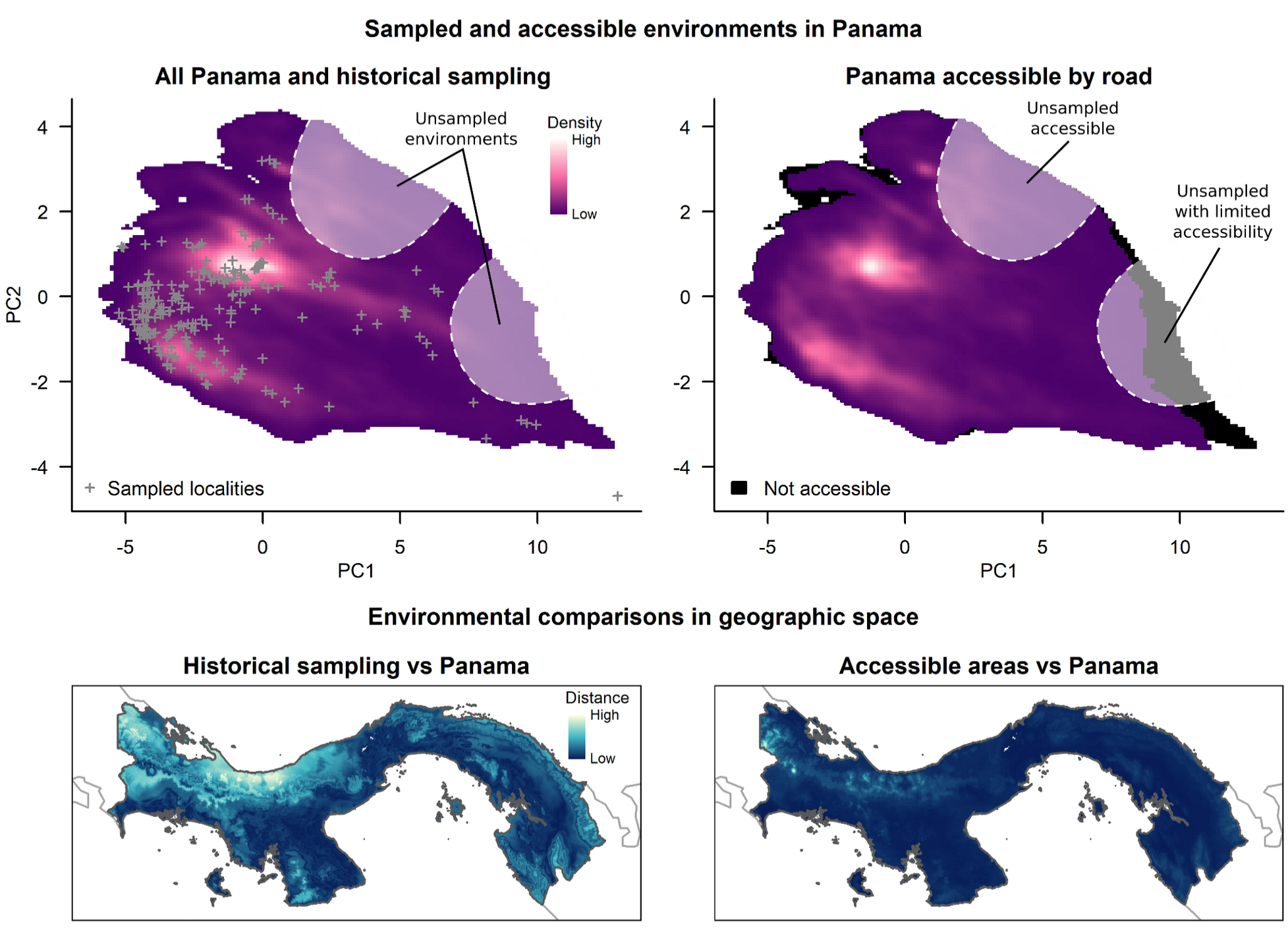
Representations of environmental space in Panama, including historically sampled (+) and accessible environments (top row). Areas of higher density in environmental space indicate conditions that are more common in the country. Historically sampled and accessible environmental conditions are compared to those present throughout the country and visualized in geographic space (bottom row). Distance refers to environmental dissimilarities between the reference conditions (i.e., sampling localities or accessible areas) and all conditions present in Panamá.

### Sites selected for pathogen surveillance

Six core sites were selected for surveillance in our initial selection process that considered criteria related to historical sampling, sampling effort, accessibility, environmental coverage, geographic representation, proximity to hantavirus outbreak sites, and distance to human settlements (Table 1, Fig. 4). These six sites are evenly distributed across available environmental conditions in Panamá and span over almost the entire geographic extent of the country. The geographic area between central Panamá and the Darién Province in the east is not represented by these six points because the environmental conditions in that area are relatively homogeneous and intermediate considering other sites selected (Fig. 4).

**Figure 4.**
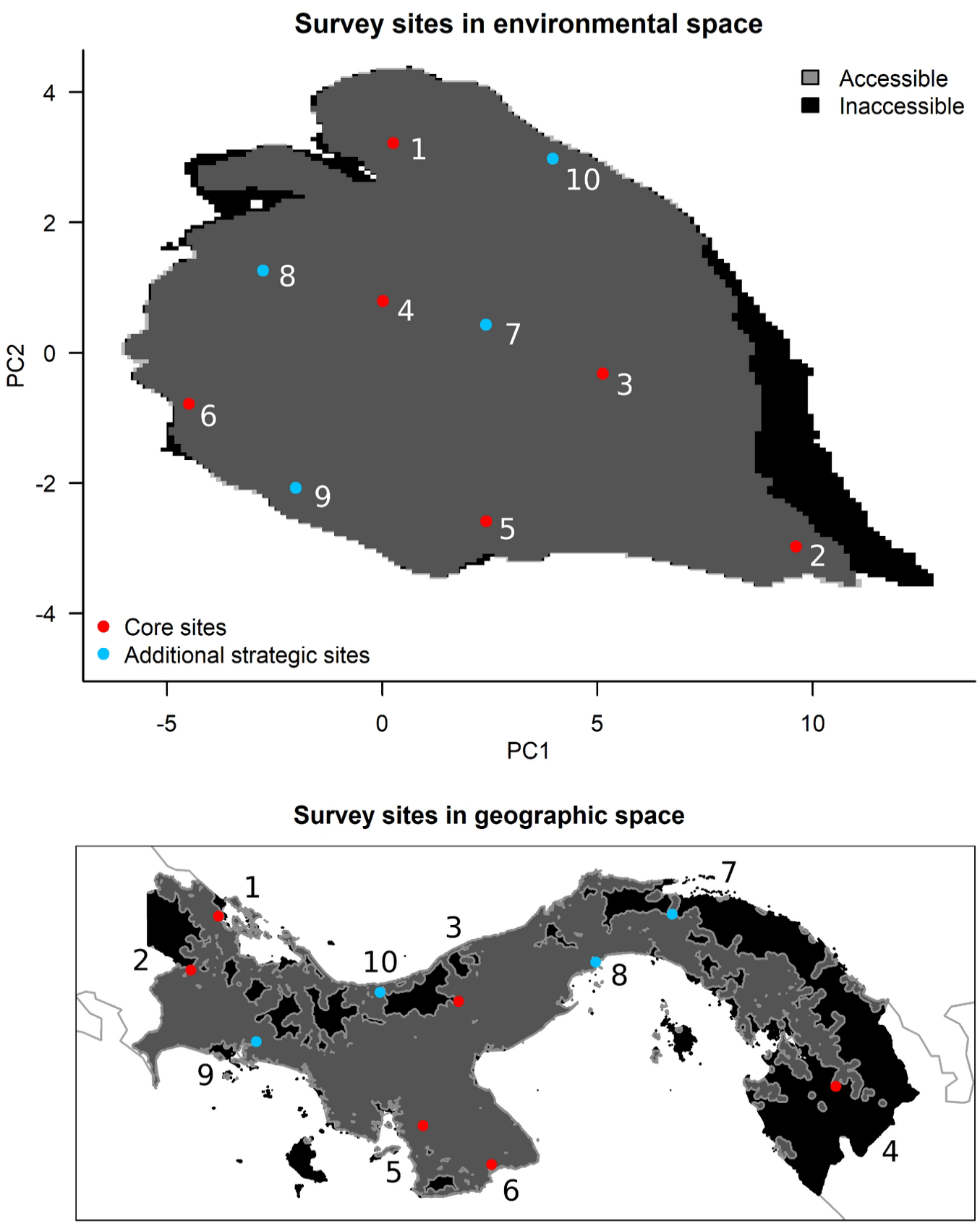
Sites selected for strategic surveillance of mammal hosts and zoonotic pathogens in Panamá in environmental (top row) and geographic space (bottom row). The six red points represent core resurvey localities and the four blue points represent additional sites of strategic significance due to environmental distinctiveness, geographic isolation, or proximity to human settlements or outbreak sites.

**Table 1.**
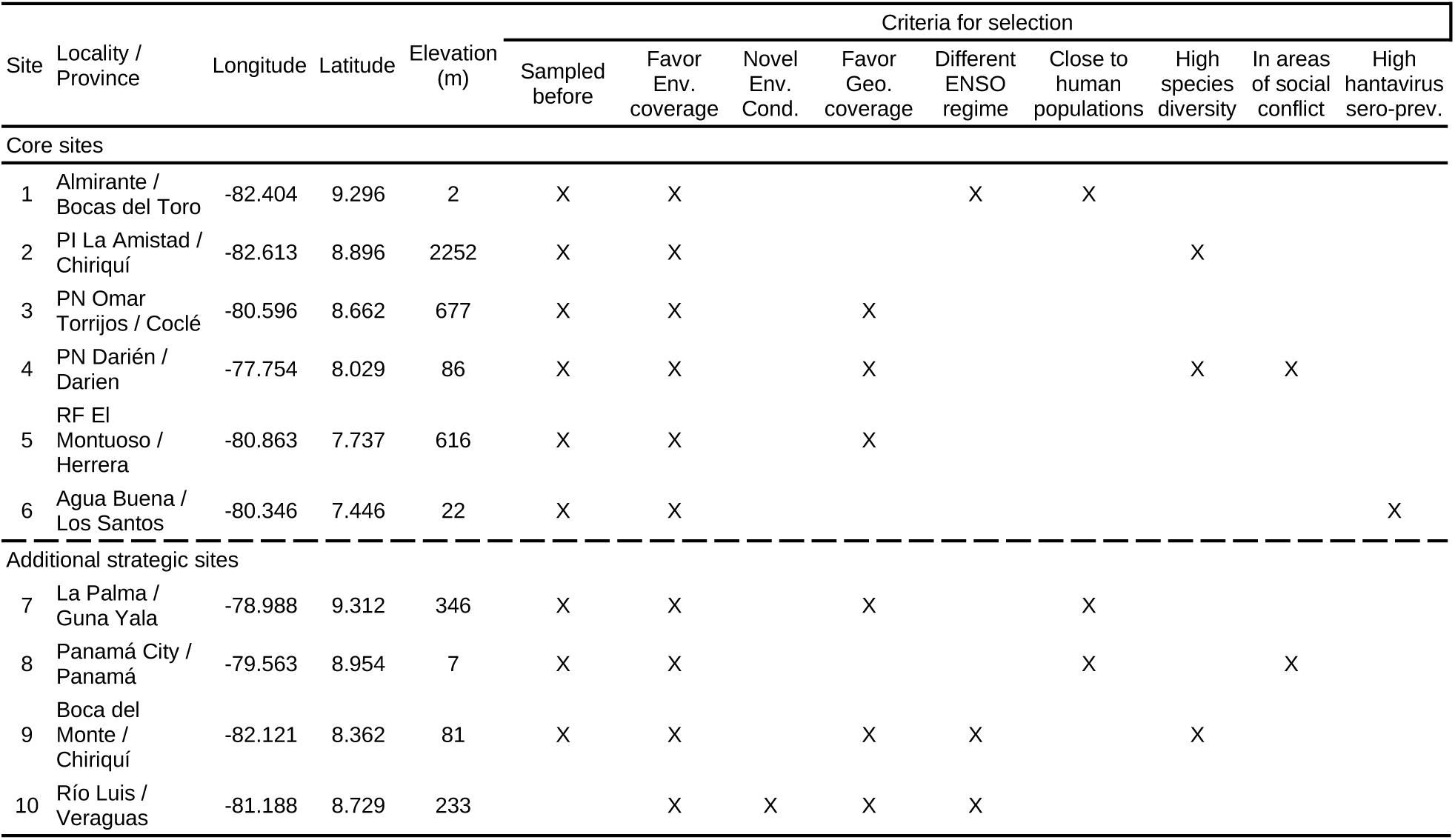
Proposed strategic surveillance sites and criteria for selection in no particular order. PI = *Parque Internacional*; PN = *Parque Nacional*; RF = *Reserva Forestal*; Env. = Environmental; Cond. = Conditions; ENSO = El Niño Southern Oscillation; Geo. = Geographic; Sero-prev. = seroprevalence in either wildlife or humans (Armién et al. 2023).

| Site | Locality / Province | Longitude | Latitude | Elevation (m) | Criteria for selection |  |  |  |  |  |  |  |  |
| --- | --- | --- | --- | --- | --- | --- | --- | --- | --- | --- | --- | --- | --- |
|  |  |  |  |  | Sampled before | Favor Env. coverage | Novel Env. Cond. | Favor Geo. coverage | Different ENSO regime | Close to human populations | High species diversity | In areas of social conflict | High hantavirus sero-prev. |
| Core sites |  |  |  |  |  |  |  |  |  |  |  |  |  |
| 1 | Almirante / Bocas del Toro | -82.404 | 9.296 | 2 | X | X |  |  | X | X |  |  |  |
| 2 | PI La Amistad / Chiriquí | -82.613 | 8.896 | 2252 | X | X |  |  |  |  | X |  |  |
| 3 | PN Omar Torrijos / Coclé | -80.596 | 8.662 | 677 | X | X |  | X |  |  |  |  |  |
| 4 | PN Darién / Darien | -77.754 | 8.029 | 86 | X | X |  | X |  |  | X | X |  |
| 5 | RF El Montuoso / Herrera | -80.863 | 7.737 | 616 | X | X |  | X |  |  |  |  |  |
| 6 | Agua Buena / Los Santos | -80.346 | 7.446 | 22 | X | X |  |  |  |  |  |  | X |
| Additional strategic sites |  |  |  |  |  |  |  |  |  |  |  |  |  |
| 7 | La Palma / Guna Yala | -78.988 | 9.312 | 346 | X | X |  | X |  | X |  |  |  |
| 8 | Panamá City / Panamá | -79.563 | 8.954 | 7 | X | X |  |  |  | X |  | X |  |
| 9 | Boca del Monte / Chiriquí | -82.121 | 8.362 | 81 | X | X |  | X | X |  | X |  |  |
| 10 | Río Luis / Veraguas | -81.188 | 8.729 | 233 |  | X | X | X | X |  |  |  |  |

We supplemented core localities, with four additional surveillance sites to: (1) help cover underrepresented geographic areas in Panamá, between the center of the country and the Darién region, that are of biogeographic relevance; (2) help complete coverage of environments in Panamá; and, perhaps more importantly, (3) sample environmental conditions never sampled before in the country, in accessible areas (site number 10; Table 1, Fig. 4). Three of these four sites have been sampled before; only one site (number 10) corresponds to a locality that has never before been sampled, but which complements representation of environments in Panama.

Our full set of localities ranges in elevation 2–2252 m and is located in 9 of the 13 provinces of Panamá. These sites represent distinct environmental conditions and the breadth of species diversity in the country, while considering historical sampling effort, distance from human settlements, and land use (some areas are agricultural lands and others are inside protected areas). Sites four and eight are of special interest because of their association with geographic areas where socio-economical conflicts are likely to occur. Site four is in a region of high interest due to human migration, and site eight is highlighted for its relationship with the Panama Canal.

## Discussion

Zoonotic pathogen surveillance is a logistically and financially challenging endeavor but one that is significantly less expensive than outbreak response and which is critical to proactive understanding of pathogen dynamics and, ultimately, risk. Zoonotic pathogens are intrinsically associated with wildlife, therefore, surveying and monitoring their wild hosts is essential to identifying public health threats prior to their emergence in humans. This case study evaluating the comprehensiveness of long-term hantavirus surveillance in Panamá outlines good practices in the design of pathogen surveillance initiatives and opportunities for refinement of existing sampling strategies. We identify a hierarchical series of strategic sampling localities in Panamá that aim to both discover novel pathogens in undersampled taxa and environments, and also refine understanding of host-pathogen ecology for known pathogens (e.g., Choclo, Calabazo hantaviruses). Selected sites build upon the network of historically sampled sites to maximize geographic and environmental dispersion and balance site accessibility and proximity to outbreak sites and human settlement. We explicitly considered environmental and geographic factors relevant to the distributions and population dynamics of pathogens, vectors, and hosts. Pathogen presence and prevalence in a host presents an environmental signal (i.e., certain conditions favor the pathogen more than others), which supports the idea of sampling broadly in environmental space. Leveraging historical sampling information allowed us to explore a set of site options that are more likely to be accessible in the future. Perhaps more importantly, considering previously sampled localities, allows for temporal comparisons that are key to characterizing changes in host-pathogen relationships over time. By considering site accessibility, we ensure that surveys can be feasibly performed. Finally, consideration of the proximity of surveillance localities to human settlements and outbreak sites makes our site selection directly relevant to public health, in addition to meaningfully contributing to permanent biodiversity infrastructure.

### Environmental representation in sampling

Historical hantavirus surveillance in Panamá has been largely reactive to existing public health threats and, consequently, biased towards particular localities (e.g., near Las Tablas). However, a deliberate effort to sample distinct geographic areas and environments (e.g., ecoregions) has substantially expanded these surveys. In fact, our results showed that the majority of accessible environments in Panamá were sampled as part of the last two decades of host-pathogen surveillance (Fig. 3). Although there is a strong bias in sampling towards certain environments (that is, the most common ones), few subsets of accessible conditions are not well sampled. Our analyses also showed that accessible areas in Panamá (within 2.5 km of a road) represent the majority of environmental combinations in the country. Environments not represented in accessible areas were rare in the country and mostly situated at high elevations, inside protected zones that are rarely visited by humans, and perhaps less likely to be implicated in spillover events. As a result, a set of survey localities that comprehensively samples environments in accessible areas is likely to cover the breadth of environments available in the country.

Thanks to the significant investment in proactive surveillance and foresight to preserve collected material and data in perpetuity with natural history museums, we only recommend adding a single new locality (Río Luis, Veraguas) to cover undersampled, yet accessible environments. Sampling that novel environment would augment the comprehensiveness of continued in-country monitoring efforts. Additional information derived from historical, boots-on-the-ground sampling directly informed site selection because experience working in those localities already exists and can facilitate the planning of a survey strategy, because logistical factors like time, habitat, local regulations, and degree of difficulty are known.

### The role of natural history collections

Recent explorations suggest that a minimum sample size of 218 is required to detect a pathogen at 15% prevalence in a host (n = Z^2^*P*[1 - *P*] / d^2^; Daniel and Cross 2019; Naing et al. 2022). Hantavirus seroprevalence at historically sampled sites in Panama ranged from 0 to 20% in most cases, when aggregating the data by month (Fig. S2). Because of the large sample sizes required for pathogen research, biomedical surveillance has relied disproportionately on fast, affordable, and non-lethal screening methods (e.g., serology) to detect the presence/absence of a pathogen of interest. While those methods provide a coarse snap-shot of pathogen presence/absence across the landscape, opportunities for validation and extension of that work, and specifically those samples, is severely limited (Colella et al. 2023). Understanding disease risk as it relates to host-pathogen interactions and their demographic fluctuations in response to environmental variation, requires a long term record (longitudinal data) of host populations and their associated microorganisms. A way to realistically achieve such large sample sizes and longitudinal coverage is via multiple-year and -locality sampling efforts linked to permanent physical specimens.

Intentional preservation of tissue samples and other specimen parts at the Museum of Southwestern Biology throughout the >20 years of hantavirus surveillance in Panama has developed a permanent, openly available, and unparalleled time series of hosts and pathogens that continues to yield insights into in-country host-pathogen dynamics (Gonzalez et al. 2023). In this way, the Gorgas Memorial Institute for Health Studies’ surveillance model exemplifies the value of pairing biomedical sampling and disease screening with holistic collecting and permanent archival of specimens in a wildlife repository. Permanent archival critically enables reevaluations of host taxonomy, validation of initial pathogen test results, and detailed investigations of host or virus biology (e.g., virology, immunology, genomics, etc.) at a later date (Dunnum et al. 2017; Colella et al. 2021). PCR and sequencing of archived tissues from seropositive samples, for example, will provide better evidence of active versus past infection (Francino et al. 2006; Tabar et al. 2009; Buchan et al. 2019). That information is necessary to narrow the time window in which infection or spillover occurred, which can help determine the specific environmental conditions influencing pathogen maintenance and transmission (Colella et al. 2023).

### Towards a strategic plan for zoonotic pathogen surveillance

Here, we evaluate the efficacy of historical hantavirus surveillance in Panamá in representing the suite environmental conditions present in Panamá, and provide recommendations for strategic surveillance sites moving forward. Although our selection was done with general consideration of site accessibility, a complete plan for surveillance will require further decision making. For example, logistical requirements, the periodicity of sampling, and the number of personnel required to comprehensively survey an area are essential parts of such a plan. We provide six recommendations for the implementation of this plan. (1) Ideally, each site will be surveyed at least twice a year to match the two main weather seasons in the country (rainy and dry). If resources are limited, surveys could be done once a year trying to visit sites in different seasons in alternating years. (2) The six core localities, together with site number 10, are essential to characterize most of the environments in Panamá, and should be prioritized in survey efforts if resources are limited. (3) Sampling effort, in terms of the number of trap nights, the traps used, transect lengths, and or grid size, should be consistent among sites and visits. (4) Future surveillance efforts should expand taxonomic sampling to improve resolution into pathogen dynamics and spillover potential for a broader suite of zoonotic pathogens. (5) Strategic alliances with key stakeholders and among public and private institutions can help facilitate and cumulatively contribute to this effort (Yeh et al. 2021; Parkinson et al. 2023; Neves et al. 2023). Finally, (6) surveys should aim to holistically collect specimens (e.g., preserve all parts to maximize the scientific value of each sampling event; Galbreath et al. 2019) to grow primary biodiversity infrastructure in-country and facilitate long term research efforts. Of course, practical decisions will have to be made that consider the unique aspects of each locality, the breadth of host diversity to be sampled, and the resources available for surveillance. As with any surveillance or management plan, methods and sites must then be periodically reevaluated in the context of new information and modified as needed (Holtrop et al. 2010; Burton et al. 2014; Meyer et al. 2015).

### Conclusions

We provide a series of recommendations for the refinement of on-going hantavirus surveillance in Panamá and for the strategic development of zoonotic pathogen surveillance programs. Overall, 20 years of surveillance in Panamá has represented most environments present in accessible areas in the country. Resampling of historically surveyed sites through time, with the addition of a few environmentally novel localities, will well represent the breadth of available environments. Such refinement is key to the effectiveness of long-term surveillance programs, which must be adaptively updated in the face of new information. Our site selection methods strike a balance between optimal environmental dispersion, site accessibility, and availability of historical samples, which provide temporal context, and can be extended to surveillance of any host-pathogen system. The strategic sites for pathogen surveillance selected have the potential to help not only in novel pathogen discovery (such as in undersampled taxa/environments), but also in refining our understanding of the reservoir/pathogen ecology and evolution of the pathogens currently known (e.g., Choclo and Calabazo hantaviruses). Last, although we have proposed a series of environmentally-optimal and accessible sites, surveillance will ultimately require significant governmental or other investment to support the personnel and logistical requirements of this work and the on-going management and preservation of resulting specimens.

## Acknowledgments

We thank the Panamanian Ministry of Environmental Affairs, the Gorgas Committee for Animal Care and Use, the Gorgas Memorial Institute for Health Studies (GMISH), the Panamanian Institute of Livestock and Agricultural Research, the Ministry of Health, the Ministry of Agricultural Development, and Smithsonian Tropical Research Institute (S. Heckadon, O. Sanjur, I. Ochoa, and A. Herre) for their support. We also thank individuals from the communities and areas surveyed, the rodent ecology team of the Ministry of Health and GMISH, especially F. Gracia, C. Muñoz, M. Ávila, E. Broce, O. Vargas, F. Crespo, C. Falcon, R. Rodriguez, J. B. Navarro, S. Bosh, R. Cedeño, D. Gonzalez, C. Falconet, J. M. Montenegro, M. Vergara, V. Dominguez, A. Perez, J. Gonzalez, A. Vergara, H. Rivera, C. Lombardo, D. Archibold, L. Castillo, A. Ortiz, D. Soto, C. Ortiz, A. Araúz, J. De Leon, J. Díaz, R. García, J. Santizo, J. F. Tello, J. Garzón, G. Niño, R. Rodríguez, D. Olivares, J. Castillo, C. Olivares, E. Salazar, P. Gutiérrez, L. Aldobán, O. Castro, J. Valencia, A. Mepaquito, S. Bethancourt, E. Ramos, and R. Figueroa for support during fieldwork. We thank the lab team Y. Zaldivar, J. Castillo, M. Ortega, M. Ramos, N. Vega, J. Correa, J. Cedeño, J. Herrera, J. Montenegro, R. Cumbrera, V. Ventura, K. Miranda, and S. Aldrette for screening thousands of samples. We thank C. Dominguez and N. Gonzalez, database managers, and R. de Vargas and I. Reyes for significant administrative support at the Department of Research in Emerging and Zoonotic Infectious Diseases, Gorgas Memorial Institute for Health Studies.

## Supplementary materials

**Table S1.** Summary of historical sampling and hantavirus seroprevalence based on serology tests.

| Species | Total | Positive serology | Hantavirus seroprevalence |
| --- | --- | --- | --- |
| *Caluromys derbianus* | 1 | 0 | 0.00 |
| *Dasypus novemcinctus* | 1 | 0 | 0.00 |
| *Didelphis marsupialis* | 10 | 0 | 0.00 |
| *Heteromys australis* | 2 | 0 | 0.00 |
| *Heteromys desmarestianus* | 65 | 0 | 0.00 |
| *Hoplomys gymnurus* | 8 | 0 | 0.00 |
| *Liomys adspersus* | 886 | 0 | 0.00 |
| Mammalia | 1 | 0 | 0.00 |
| *Marmosa* spp. | 1 | 0 | 0.00 |
| *Marmosa isthmica* | 2 | 0 | 0.00 |
| *Marmosa robinsoni* | 11 | 0 | 0.00 |
| *Melanomys caliginosus* | 40 | 0 | 0.00 |
| *Metachirus nudicaudatus* | 1 | 0 | 0.00 |
| *Mus musculus* | 463 | 2 | 0.00 |
| *Nephelomys devius* | 39 | 0 | 0.00 |
| *Nyctomys sumichrasti* | 3 | 0 | 0.00 |
| *Oecomys trinitatis* | 5 | 0 | 0.00 |
| *Oligoryzomys costaricensis* | 822 | 138 | 0.17 |
| *Oligoryzomys vegetus* | 11 | 0 | 0.00 |
| *Oryzomys* spp. | 10 | 0 | 0.00 |
| *Oryzomys couesi* | 6 | 0 | 0.00 |
| *Peromyscus nudipes* | 130 | 6 | 0.05 |
| *Proechimys semispinosus* | 163 | 0 | 0.00 |
| *Rattus norvegicus* | 3 | 0 | 0.00 |
| *Rattus rattus* | 165 | 0 | 0.00 |
| *Reithrodontomys creper* | 64 | 4 | 0.06 |
| *Reithrodontomys darienensis* | 2 | 0 | 0.00 |
| *Reithrodontomys fulvescens* | 1 | 0 | 0.00 |
| *Reithrodontomys mexicanus* | 78 | 23 | 0.29 |
| *Reithrodontomys sumichrasti* | 9 | 4 | 0.44 |
| *Sciurus granatensis* | 1 | 0 | 0.00 |
| *Scotinomys teguina* | 1 | 0 | 0.00 |
| *Scotinomys xerampelinus* | 96 | 0 | 0.00 |
| *Sigmodon hirsutus* | 1186 | 46 | 0.04 |
| *Sigmodontinae* | 4 | 0 | 0.00 |
| *Sigmodontomys alfari* | 2 | 0 | 0.00 |
| *Transandinomys bolivaris* | 29 | 0 | 0.00 |
| *Transandinomys talamancae* | 242 | 2 | 0.01 |
| *Tylomys watsoni* | 1 | 0 | 0.00 |
| *Zygodontomys brevicauda* | 3525 | 276 | 0.08 |

**Table S2.**
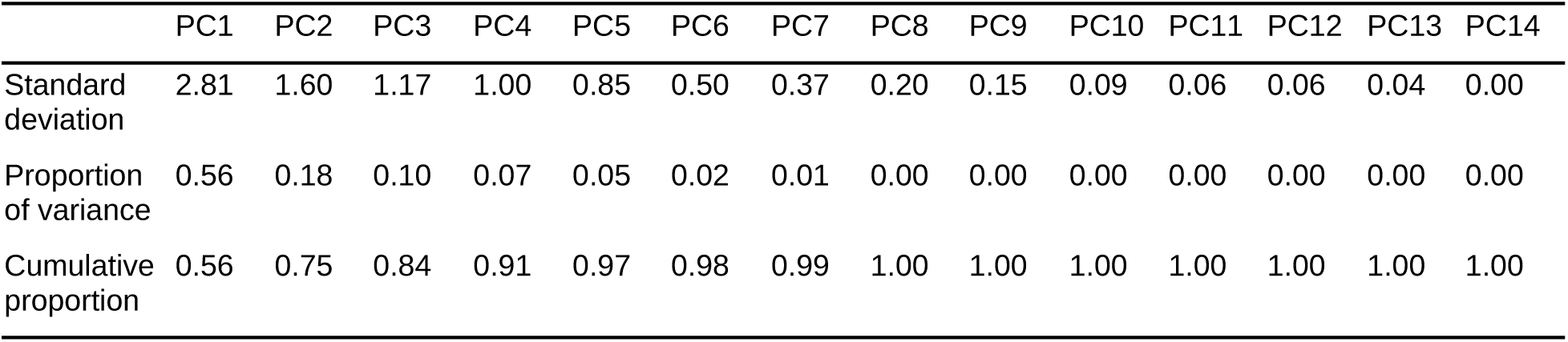
Summary of variance explained by principal components deriving from analysis of environmental variables.

**Table S3.** Principal component analysis loadings. Variables are as follows: annual mean temperature (bio1), maximum temperature of warmest month (bio5), minimum temperature of colder month (bio6), temperature annual range (bio7), annual precipitation (bio12), precipitation seasonality (bio15), precipitation of wettest quarter (bio16), precipitation of driest quarter (bio17), maximum potential evapotranspiration (pet_max), mean potential evapotranspiration (pet_mean), minimum potential evapotranspiration (pet_min), maximum vapor pressure deficit (vpd_max), mean vapor pressure deficit (vpd_mean), and minimum vapor pressure deficit (vpd_min).

|  | PC1 | PC2 | PC3 | PC4 | PC5 | PC6 | PC7 | PC8 | PC9 | PC10 | PC11 | PC12 | PC13 | PC14 |
| --- | --- | --- | --- | --- | --- | --- | --- | --- | --- | --- | --- | --- | --- | --- |
| bio1 | -0.30 | 0.21 | -0.16 | -0.23 | -0.28 | -0.24 | -0.18 | 0.13 | 0.05 | -0.14 | -0.46 | -0.46 | -0.40 | 0.00 |
| bio12 | 0.24 | -0.17 | -0.56 | 0.19 | -0.11 | -0.06 | -0.06 | 0.03 | -0.07 | 0.11 | -0.43 | -0.11 | 0.57 | 0.00 |
| bio15 | -0.20 | -0.44 | 0.19 | 0.12 | -0.35 | 0.22 | 0.16 | 0.14 | 0.69 | -0.04 | -0.11 | 0.01 | 0.09 | 0.00 |
| bio16 | 0.16 | -0.35 | -0.51 | 0.26 | -0.27 | 0.06 | -0.05 | 0.10 | -0.13 | -0.07 | 0.36 | 0.08 | -0.53 | 0.00 |
| bio17 | 0.24 | 0.29 | -0.37 | -0.02 | 0.41 | -0.26 | 0.17 | -0.06 | 0.65 | -0.06 | 0.06 | 0.06 | -0.14 | 0.00 |
| bio5 | -0.31 | -0.14 | -0.18 | -0.38 | -0.03 | -0.26 | -0.23 | 0.12 | 0.05 | 0.06 | 0.26 | 0.23 | 0.20 | -0.64 |
| bio6 | -0.27 | 0.29 | -0.15 | -0.07 | -0.44 | -0.27 | 0.00 | -0.04 | 0.04 | 0.07 | 0.29 | 0.29 | 0.23 | 0.57 |
| bio7 | -0.08 | -0.50 | -0.06 | -0.40 | 0.44 | -0.02 | -0.29 | 0.20 | 0.01 | 0.00 | 0.00 | -0.04 | -0.01 | 0.51 |
| pet_max | -0.29 | -0.24 | 0.05 | 0.26 | 0.18 | -0.52 | 0.55 | 0.13 | -0.16 | 0.33 | 0.02 | -0.17 | -0.05 | 0.00 |
| pet_mean | -0.31 | -0.04 | 0.01 | 0.43 | 0.22 | -0.18 | -0.12 | 0.03 | -0.08 | -0.72 | -0.13 | 0.26 | 0.08 | 0.00 |
| pet_min | -0.30 | 0.07 | 0.01 | 0.48 | 0.16 | 0.04 | -0.58 | -0.24 | 0.17 | 0.45 | 0.08 | -0.13 | -0.05 | 0.00 |
| vpd_max | -0.32 | -0.15 | -0.23 | -0.19 | 0.06 | 0.21 | 0.24 | -0.66 | -0.07 | 0.09 | -0.32 | 0.32 | -0.18 | 0.00 |
| vpd_mean | -0.33 | 0.08 | -0.26 | -0.05 | 0.12 | 0.36 | 0.22 | -0.12 | -0.02 | -0.24 | 0.41 | -0.57 | 0.24 | 0.00 |
| vpd_min | -0.28 | 0.30 | -0.21 | 0.07 | 0.18 | 0.44 | 0.13 | 0.61 | -0.06 | 0.22 | -0.16 | 0.29 | -0.07 | 0.00 |

**Figure S1.**
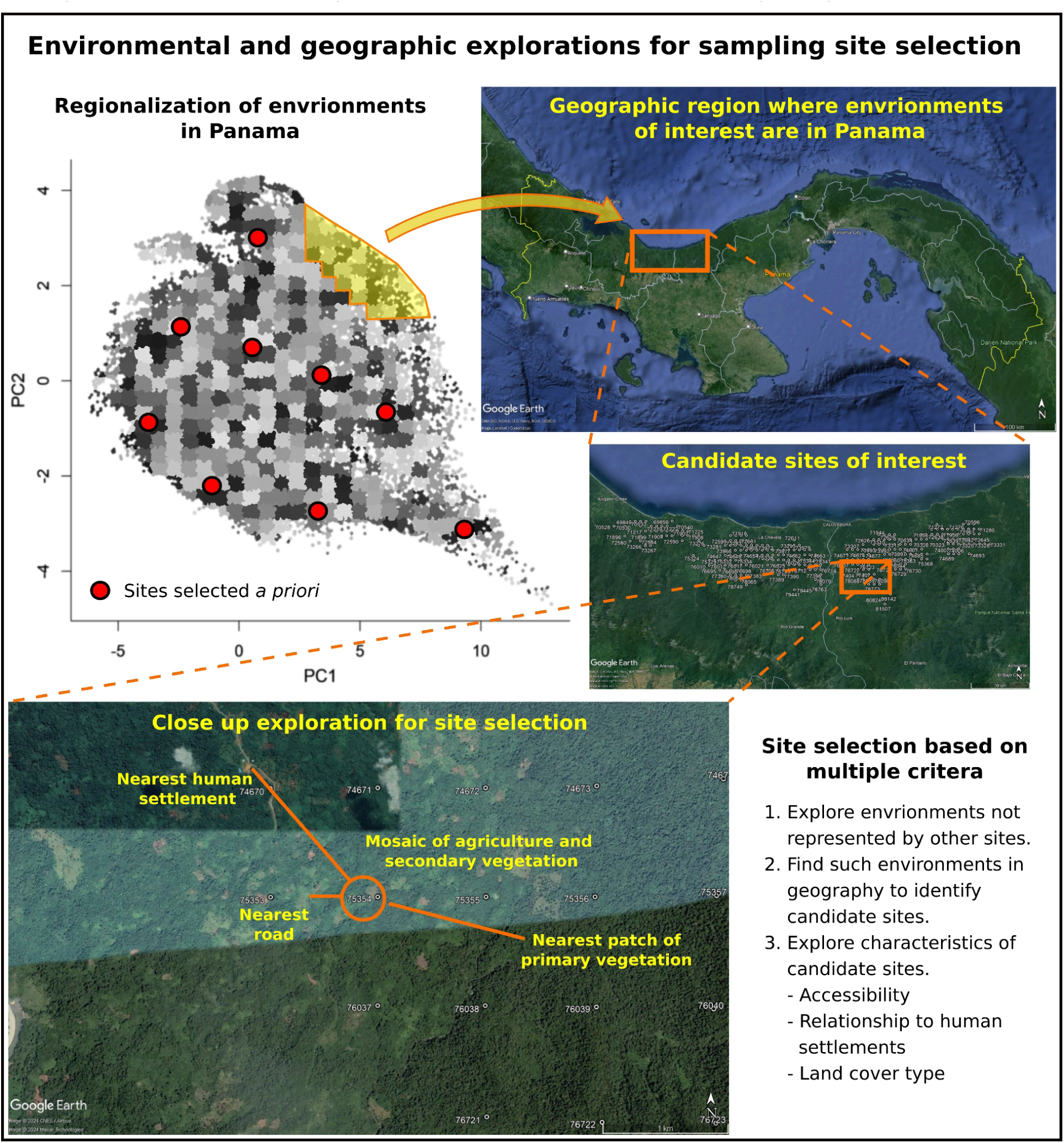
Schematic representation of the workflow for sampling site selection illustrated with a locality in an environmental region not included in localities historically sampled in Panama.

**Figure S2.**
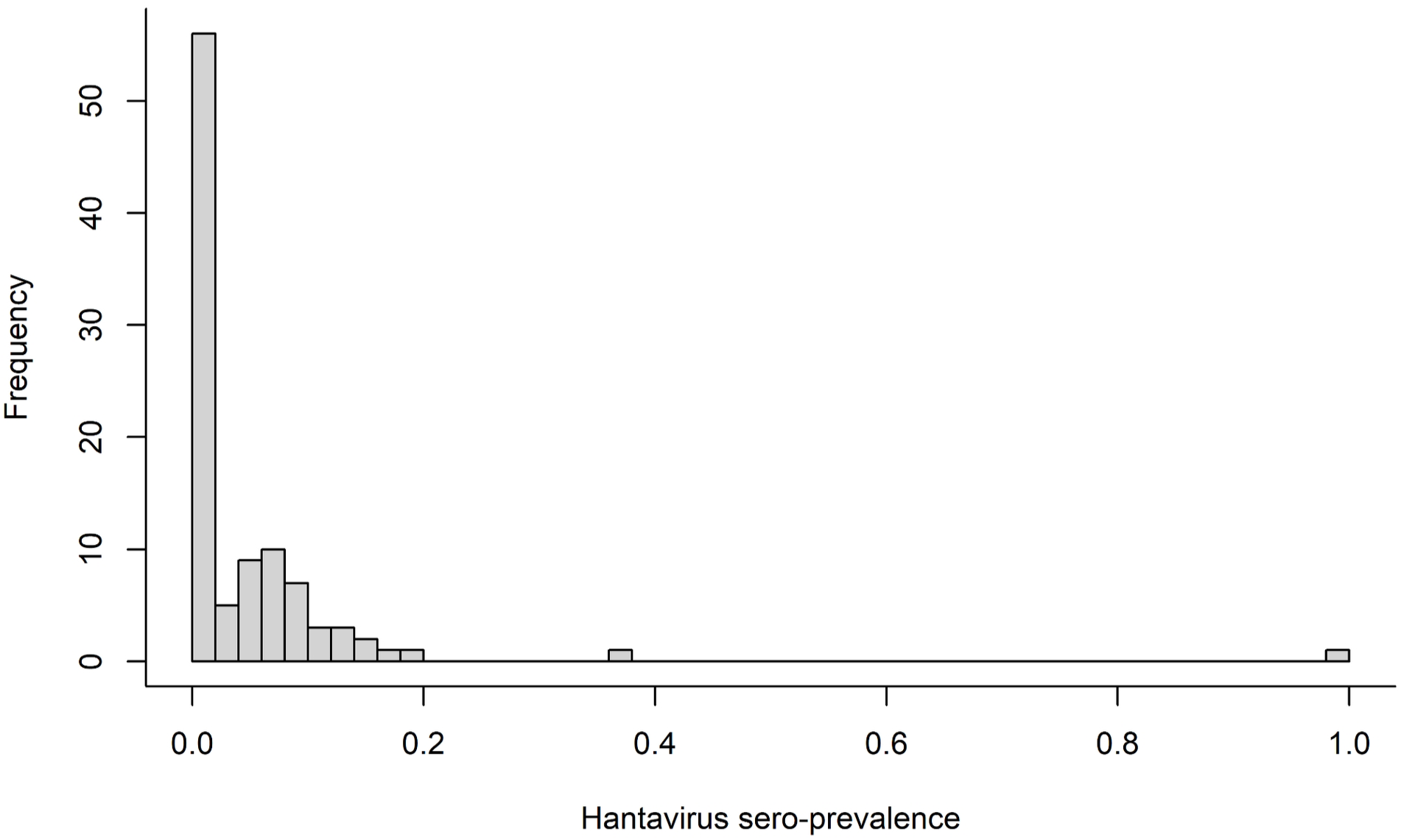
Summary of hantavirus seroprevalence in historical samples aggregated by month.

